# AAV-mediated age- and circuit-dependent restoration of photoreceptor synaptic structure and function in α2δ4-associated retinal synaptopathy

**DOI:** 10.64898/2026.07.21.739956

**Authors:** Trong Thuan Ung, Gillian N. Huskin, Sanford Boye, Yuchen Wang

## Abstract

**Purpose:** Synaptic dysfunction and neurite remodeling are early features of many inherited retinal diseases, but whether synaptic function can be restored after circuit remodeling remains unclear. We evaluated the therapeutic potential of adeno-associated virus (AAV)-mediated α2δ4 gene supplementation in a mouse model of *CACNA2D4*-associated retinal synaptopathy.

**Methods:** AAV8-GRK1-α2δ4 was delivered by subretinal injection to α2δ4 knockout mice at neonatal, young adult, or middle-aged adult stages. Retinal function was assessed by electroretinography. Synaptic organization and circuit remodeling were evaluated by immunohistochemistry.

**Results:** AAV-mediated α2δ4 expression restored synaptic localization of Cav1.4, ELFN proteins, and mGluR6 at all treatment ages. However, circuit structure and functional rescue were age- and circuit-dependent. Neonatal treatment restored rod and cone transmission and prevented photoreceptor terminal retraction and bipolar cell dendritic sprouting. Rescue in young adults improved rod and cone circuit function but did not reverse neurite remodeling. Rescue in middle-aged mice restored cone but not rod transmission. Notably, neonatal treatment produced the highest proportion of synapses within synaptic layer, whereas adult treatment generated increasing ectopic synapses in nuclear layer.

**Conclusions:** Photoreceptor synaptic assembly remains plastic in remodeled retinas and can be restored by α2δ4 supplementation. However, rescue efficacy is influenced by age and circuits with rod circuits exhibiting a narrower therapeutic window. Synaptic molecular reassembly can occur without neurite restoration. These findings establish α2δ4 gene therapy as a promising treatment for *CACNA2D4*-associated retinal synaptopathy and reveal distinct intervention windows for synaptic assembly, neurite remodeling, and rod versus cone circuit recovery.

## Introduction

The first visual synapses, formed between photoreceptors and bipolar cells in the outer plexiform layer, are the obligatory entry points through which rod- and cone-mediated signals are transmitted to downstream retinal circuits and the brain ^1^. Rod photoreceptors primarily synapse with rod bipolar cells to support dim-light vision, whereas cone photoreceptors contact multiple ON and OFF cone bipolar cell types to encode daylight and color vision ^2–4^. Because all visual information must pass through these highly specialized ribbon synapses, defects in their assembly, maintenance, or spatial organization can profoundly compromise retinal signaling ^5^.

Synaptic dysfunction at the first visual synapses is a central feature of congenital stationary night blindness and is increasingly recognized as an early event in inherited retinal degenerations, including retinitis pigmentosa and cone-rod dystrophy ^6–9^. In these conditions, disruption of synaptic molecular organization often precedes or accompanies structural remodeling, including photoreceptor terminal retraction and bipolar cell dendritic sprouting ^5^. Thus, the photoreceptor-to-bipolar cell synapse is not only a critical site of early disease, but also a potential therapeutic target for preserving or restoring visual function.

Rod and cone terminals are ribbon-type synapses specialized for graded, sustained glutamate release ^10^. At the presynaptic active zone, the L-type voltage-gated calcium channel (VGCC) Cav1.4 couples changes in photoreceptor membrane potential to calcium influx and vesicle release ^11–13^. On the postsynaptic side, rod bipolar cells and ON cone bipolar cells concentrate the metabotropic glutamate receptor mGluR6 at dendritic tips, where it initiates the ON pathway response ^14–16^.

A key molecule linking the Cav1.4-containing release apparatus to the postsynaptic mGluR6 receptor complex is α2δ4, an auxiliary subunit of voltage-gated calcium channels, encoded by *CACNA2D4* ^17,18^. α2δ4 regulates Cav1.4 abundance, localization, and function ^19–22^. In addition, α2δ4 is required for proper localization of trans-synaptic cell adhesion molecules, including ELFN1 at rod synapses and ELFN2 at cone synapses, which directly interact with postsynaptic mGluR6 and control its dendritic targeting ^23,24^. Loss of α2δ4 disrupts the molecular architecture of both rod and cone synapses, causing mislocalization of Cav1.4, ELFN proteins, and mGluR6. Moreover, α2δ4 loss also induces structural remodeling, including rod terminal retraction, cone pedicle shrinkage, and bipolar cell dendritic sprouting ^20,21^, phenotypes that resemble synaptic remodeling observed in multiple retinal diseases ^5^.

These studies provide mechanistic insight into night blindness and cone dysfunction associated with *CACNA2D4* mutations ^25,26^. However, whether gene supplementation can restore synaptic structure and function after α2δ4 loss remains incompletely understood. Previous neonatal rescue experiments demonstrated recovery of selected rod synaptic markers, including ELFN1 and mGluR6, but did not determine whether functional transmission was restored ^21^. Moreover, because human *CACNA2D4*-associated disease affects both rod and cone pathways and may not be clinically recognized until later in life ^25,26^, it is important to define whether AAV-mediated α2δ4 supplementation can rescue both circuits across developmental and adult stages. Such studies are essential for establishing the therapeutic window for α2δ4 gene therapy and for identifying which aspects of synaptic remodeling remain reversible in the mature diseased retina.

Here, we performed AAV-mediated α2δ4 rescue in α2δ4-deficient mice at neonatal, young adult, and middle-aged adult stages, followed by quantitative analyses of retinal function and synaptic organization. We found that α2δ4 supplementation restored synaptic molecular organization and improved function in both developing and mature knockout retinas. However, rescue outcome was strongly shaped by age and circuit identity. Neonatal rescue restored both synaptic organization and neurite elaboration, whereas adult rescue reassembled synaptic molecular complexes without correcting ectopic neurite remodeling. In addition, rod circuit function was more sensitive to the age of intervention than cone circuit function, indicating that cone synapses retain a broader therapeutic window. Together, these findings reveal that synaptic molecular organization and neurite positioning are governed by separable mechanisms with distinct developmental constraints, and they provide a framework for developing α2δ4-targeted gene therapy for inherited retinal synaptopathies.

## Results

### Neonatal α2δ4 rescue restores synaptic structure and function in both rod and cone circuits

To restore α2δ4 expression in photoreceptors, we generated an AAV8 vector expressing full-length mouse α2δ4 under the control of the GRK1 promoter, which drives expression in both rods and cones ^27–30^. We first tested this strategy in neonatal α2δ4 KO mice by delivering AAV8-GRK1-α2δ4 subretinally to one eye and a control AAV8-GRK1-GFP vector to the contralateral eye. Retinal function and synaptic structure were examined one month after injection using electroretinography (ERG) and immunohistochemistry (IHC) (Fig. 1A).

**Figure 1.**
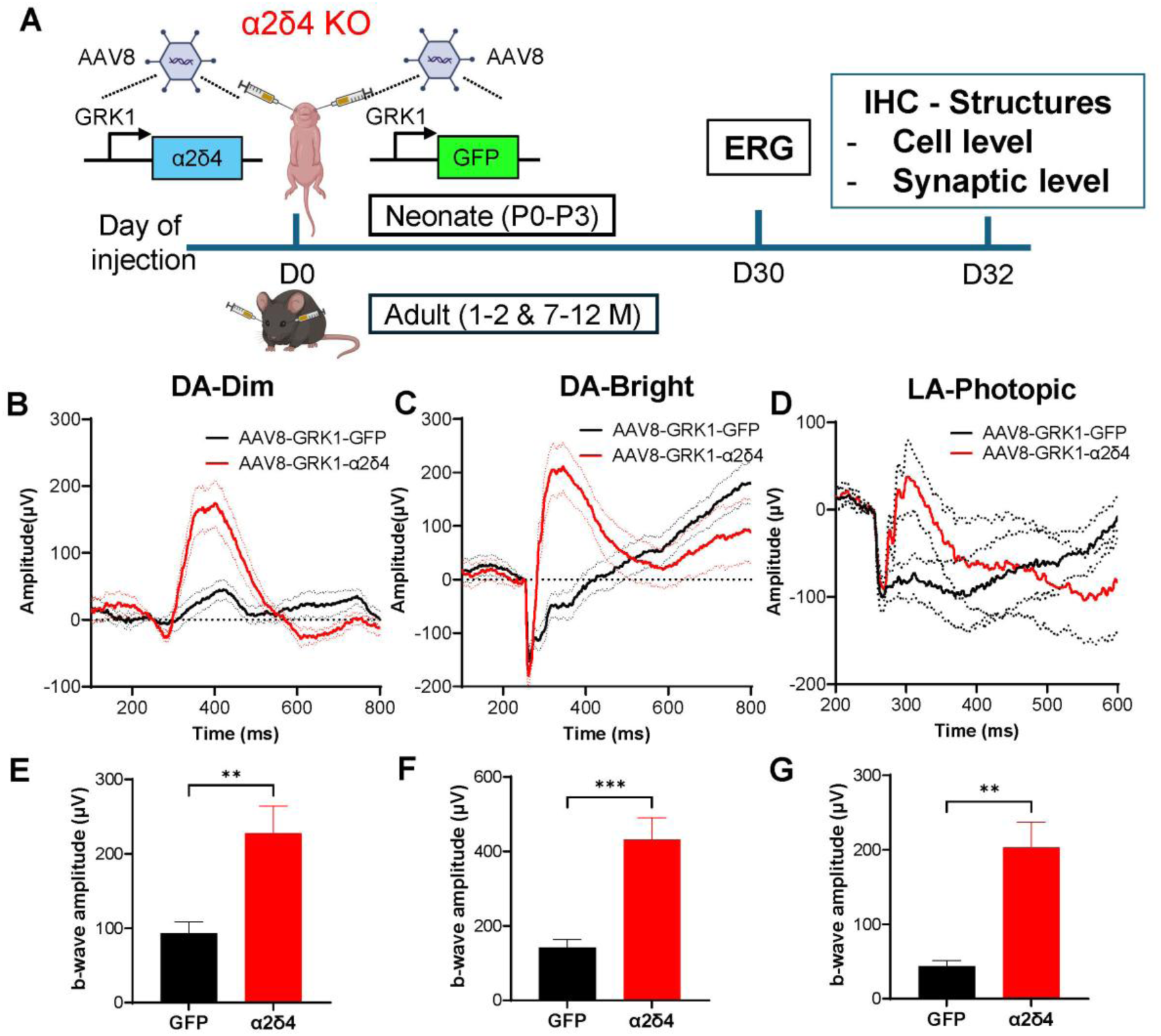
Electroretinography (ERG) studies demonstrate that rescue in α2δ4 KO neonates restores synaptic function in both rod and cone circuits. **A.** Schematic of the rescue study design showing subretinal injection of GRK1 promoted α2δ4-expressing AAV8 to one eye and control GFP AAV to the other in α2δ4 KO mouse followed by ERG and IHC studies at day 30 after injection. **B-C.** Average traces of dark-adapted ERG response to dim light 2.479 log photons/µm^2^ (**B**) which primarily activates rods, and bright light 6.316 log photons/µm^2^ (**C**), which stimulated both rods and cones. **D.** Average traces of light-adapted ERG response to light intensity of 6.316 log photons/µm^2^ isolating cones responses from α2δ4 KO retinas. **E-F.** Quantification of b-wave amplitudes of dark-adapted ERG responses at light intensities of 2.479 (**E**) and 6.316 (**F**) log photons/µm², in rescued α2δ4 KO mice. (n = 11 mice). **G**. Quantification of b-wave amplitudes of light-adapted ERG responses at light intensities of 6.316 log photons/µm^2^ (n=9 mice). Error bars are SEM, unpaired t-test (**F**). *\*p < 0.05; **p < 0.01, ***p < 0.001; points without an asterisk are not significant (ns)*.

We assessed rod- and cone-mediated visual signaling using dark- and light-adapted ERG recordings. Under dark-adapted conditions, α2δ4-rescued eyes showed a significant increase in b-wave amplitude at both dim and bright flash intensities which were about half of the WT controls (Fig. 1B, C, E, F; Suppl. Fig. 1A-D), whereas a-wave amplitudes were unchanged compared with control eyes (Suppl. Fig. 1E). Under light-adapted conditions, which isolate cone pathway responses by saturating rods, rescued eyes also displayed a significant increase in b-wave amplitude without a detectable change in the a-wave (Fig. 1D, G; Suppl. Fig. 1F). These results indicate that neonatal α2δ4 supplementation improves synaptic transmission in both rod and cone pathways without altering photoreceptor outer retinal responses.

We next determined whether functional recovery was accompanied by restoration of synaptic architecture. We examined photoreceptor terminals and synaptic organization using markers for axonal terminals (PSD95 and cone arrestin), presynaptic ribbons (CTBP2), calcium channels (Cav1.4), trans-synaptic adhesion molecules (ELFN1 and ELFN2), and the postsynaptic receptor mGluR6, based on previously defined α2δ4 KO phenotypes ^21^. Viral transduction was confirmed by GFP expression in the outer nuclear layer, and α2δ4 protein localized appropriately to both rod and cone synapses, where it overlapped with synaptic ribbon labeling (Suppl. Fig. 2A, B). In control retinas, PSD95 labeling in the OPL was reduced and disorganized, with ectopic PSD95-positive structures extending into the ONL. In contrast, α2δ4-rescued retinas showed restored PSD95 intensity and morphology in the OPL, with no obvious ectopic PSD95-positive structures in the ONL (Fig. 2A). Cone terminals, which appeared small and faint in control retinas, also recovered in both size and labeling intensity after α2δ4 rescue (Fig. 2B). In addition, rod bipolar cell dendrites, which showed prominent sprouting into the ONL in control retinas, were largely restored to their normal OPL localization in rescued retinas (Fig. 2C).

**Figure 2.**
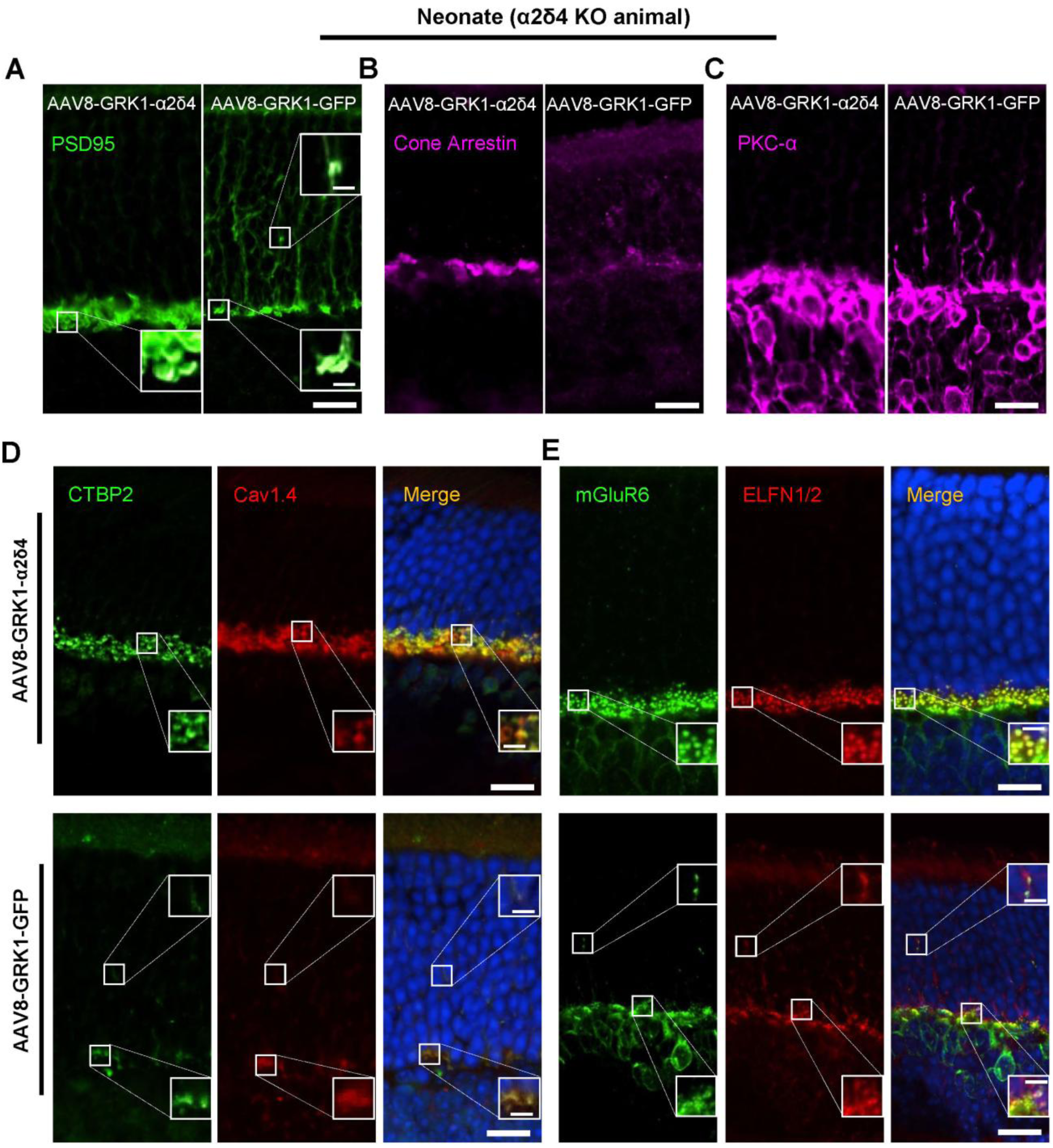
Immunohistochemistry (IHC) studies demonstrate that rescue in α2δ4 KO neonates restores synaptic structure in both rod and cone circuits. **A-C.** Representative confocal images of retinal sections from α2δ4 KO retinas stained with anti-PSD95 (**A**), anti-cone arrestin (**B**), and anti-PKCα (**C**) to label photoreceptor terminals, cone photoreceptors, and rod bipolar cells (RBCs), respectively. Scale bar, 10 μm. The insets show higher-magnification images; Scale bar, 2.5 μm. **D**. Representative confocal images of retinal sections from α2δ4 KO retinas double immunolabeled for CTBP2 (green) and Cav1.4 (red). Scale bar, 10 μm. Inset; 2.5 μm **E**. Representative confocal images of retinal sections from α2δ4 KO neonates double immunolabeled for mGluR6 (green) and ELFN1/2 (red). Scale bar, 10 μm. Inset; 2.5 μm

At the presynaptic active zone, loss of α2δ4 caused distorted ribbon labeling and markedly reduced synaptic Cav1.4 in both rod and cone terminals ^20,21^. AAV-mediated α2δ4 expression restored CTBP2 and Cav1.4 labeling in the OPL (Fig. 2D). In control KO retinas, rod-specific ELFN1 targeting was nearly abolished and cone-associated ELFN2 targeting was severely reduced, whereas both ELFN1 and ELFN2 were robustly localized to the OPL after rescue (Fig. 2E). Consistent with the role of ELFNs in organizing postsynaptic mGluR6, rescued retinas displayed prominent mGluR6 puncta at both rod and cone synapses, where they colocalized with ELFN-positive synaptic sites (Fig. 2E).

Together, these functional and structural analyses show that AAV-mediated α2δ4 expression during retinal development is sufficient to restore photoreceptor synaptic architecture, suppress neurite remodeling, and recover synaptic transmission in both rod and cone circuits.

### Young adult α2δ4 rescue restores synaptic molecular organization and function but does not reverse neurite remodeling

To determine whether mature α2δ4-deficient retinas retain the capacity for rescue, we applied the same treatment paradigm to 1- to 2-month-old α2δ4 KO mice. Dark-adapted ERG recordings showed that α2δ4-rescued eyes had significantly larger b-wave amplitudes across the tested flash intensities (Fig. 3A, B, D, E; Suppl. Fig. 3 A-C), while a-wave amplitudes remained comparable to control eyes (Suppl. Fig. 3E). Light-adapted ERG recordings also showed robust b-wave recovery without a corresponding change in the a-wave (Fig. 3C, F; Suppl. Fig. 3F). Thus, α2δ4 supplementation in young adult retinas improves synaptic transmission within both rod and cone pathways while preserving comparable photoreceptor-driven a-wave responses.

**Figure 3.**
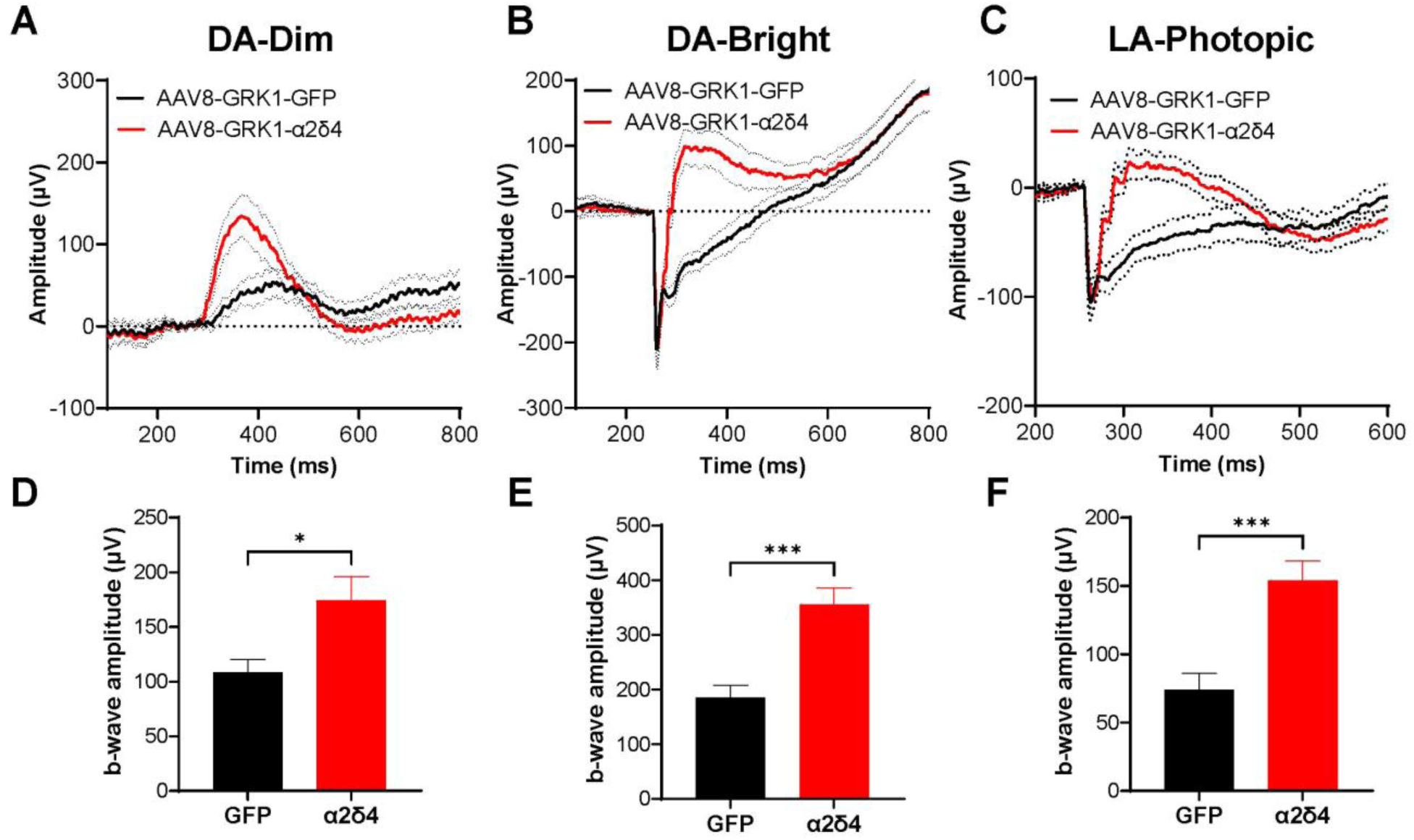
Electroretinography (ERG) studies demonstrate that rescue in young adult α2δ4 KO (1-2M) restores synaptic function in both rod and cone circuits. **A-B.** Average traces of dark-adapted ERG response to light intensity of 2.479 (**A**), 6.316 (**B**) log photons/µm^2^, from α2δ4 KO mice. **C.** Average traces of light-adapted ERG response to light intensity of 6.316 log photons/µm^2^, from α2δ4 KO mice. **D-E.** Quantification of b-wave amplitudes of dark-adapted ERG responses at light intensities of 2.479 (**D**) and 6.316 (**E**) log photons/µm², in rescued α2δ4 KO mice. (n = 13 mice). **F.** Quantification of b-wave amplitudes of light-adapted ERG responses at light intensities of 6.316 log photons/µm^2^ in rescued α2δ4 KO mice. (n = 12 mice). Error bars are SEM, unpaired t-test. *\*p < 0.05; **p < 0.01, ***p < 0.001; points without an asterisk are not significant (ns)*.

Functional recovery in young adult retinas was accompanied by substantial restoration of synaptic molecular organization. In rescued eyes, PSD95 labeling recovered in intensity and morphology within the OPL, similar to the neonatal rescue group. However, in contrast to neonatal rescue, prominent PSD95-positive structures remained in the ONL (Fig. 4A). These ectopic PSD95-positive structures likely correspond to retracted rod terminals that were not repositioned by α2δ4 supplementation. Similar ectopic localization was observed for ribbon markers, ELFNs, and mGluR6, although these synaptic puncta displayed organized, WT-like morphology (Fig. 4B-E). Quantification of ELFN and mGluR6 puncta size at both rod and cone synapses showed no detectable difference between neonatal and young adult rescue groups (Suppl. Fig. 4A-D), indicating that mature retinas can reassemble synaptic molecular complexes even when neurite positioning remains abnormal.

**Figure 4.**
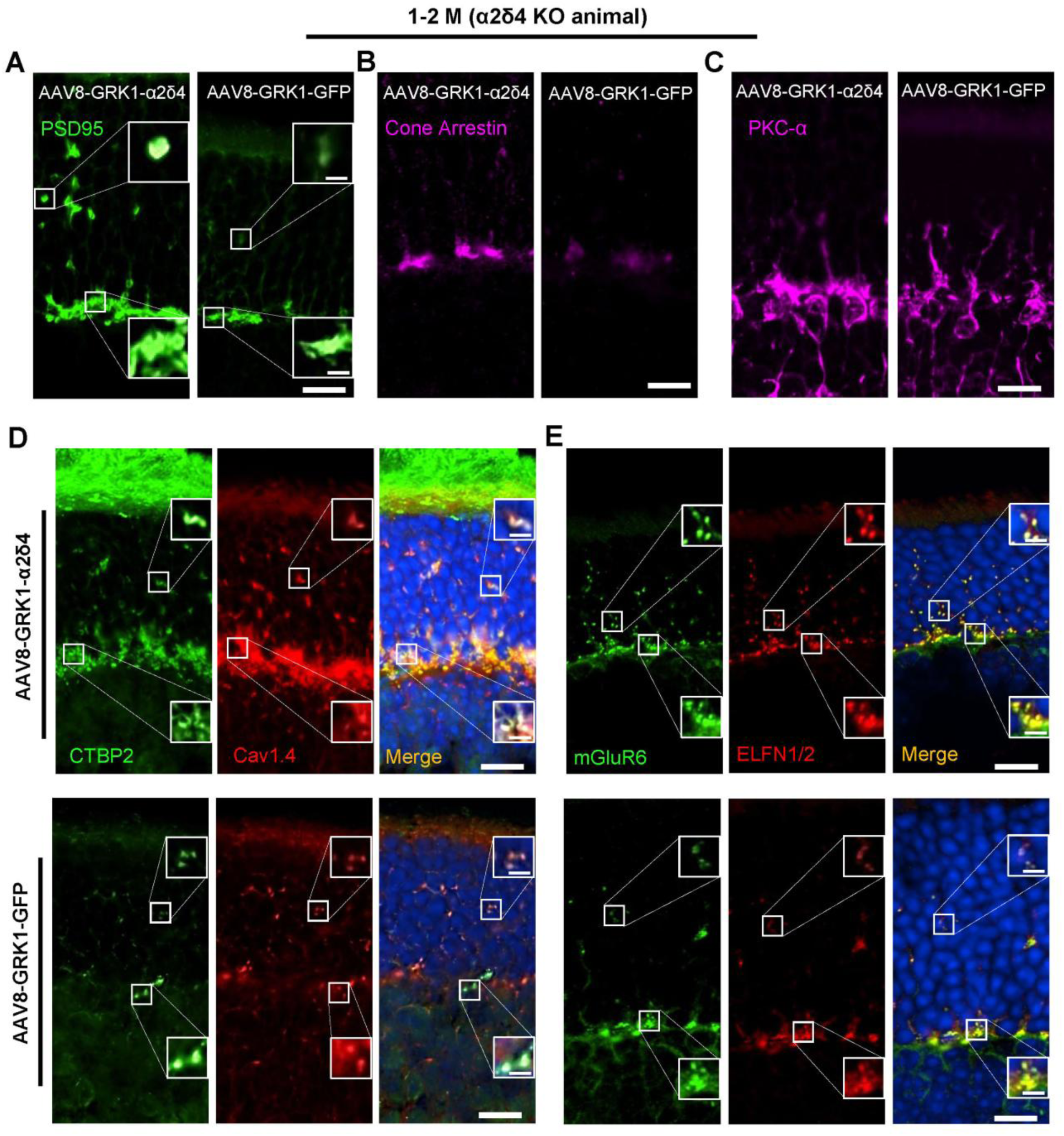
IHC studies demonstrate that rescue in young adult α2δ4 KO (1-2M) restores synaptic assembly but not neurite remodeling. **A-C.** Representative confocal images of retinal sections from α2δ4 KO mice stained with anti-PSD95 (**A**), anti-cone arrestin (**B**), and anti-PKCα (**C**) to label photoreceptor terminals, cone photoreceptors, and rod bipolar cells (RBCs), respectively. Scale bar, 10 μm. The inset shows a higher-magnification image; Scale bar, 2.5 μm. **D**. Representative confocal images of retinal sections from α2δ4 KO mice double immunolabeled for CTBP2 (green) and Cav1.4 (red). Scale bar, 10 μm. Inset, 2.5 μm. **E**. Representative confocal images of retinal sections from α2δ4 KO mice double immunolabeled for mGluR6 (green) and ELFN1/2 (red). Scale bar, 10 μm. Inset, 2.5 μm.

These findings demonstrate that young adult α2δ4 KO retinas remain competent for molecular and functional rescue. However, the persistence of ectopic synaptic structures in the ONL suggests that α2δ4-dependent synaptic molecular assembly and α2δ4-associated neurite remodeling are governed by distinct mechanisms, with neurite positioning showing a stronger dependence on developmental timing.

### Middle-aged adult α2δ4 rescue preserves cone pathway recovery but shows reduced rod pathway rescue

Because visual impairment associated with *CACNA2D4* mutations may not be recognized until adulthood ^25,26^, we tested whether α2δ4 gene supplementation remains effective in older α2δ4 KO mice. We performed subretinal rescue in 7- to 12-month-old mice, an age range corresponding to middle-aged adulthood in humans. Dark-adapted ERG recordings showed a significant increase in b-wave amplitude at bright flash intensities, where both rod and cone pathways contribute, while a-wave amplitudes remained comparable between rescued and control eyes (Fig. 5B, D; Suppl. Fig. 5A). However, at dim scotopic flash intensities that preferentially assess rod pathway function, b-wave amplitudes were not significantly increased in rescued eyes compared with controls (Fig. 5A, D). In contrast, light-adapted ERG recordings, which isolate cone pathway function, showed a significant increase in b-wave amplitude without a change in the a-wave (Fig. 5C, E; Suppl. Fig. 5B). These results suggest that cone pathway rescue remains robust in middle-aged retinas, whereas rod pathway functional recovery becomes more limited with age.

**Figure 5.**
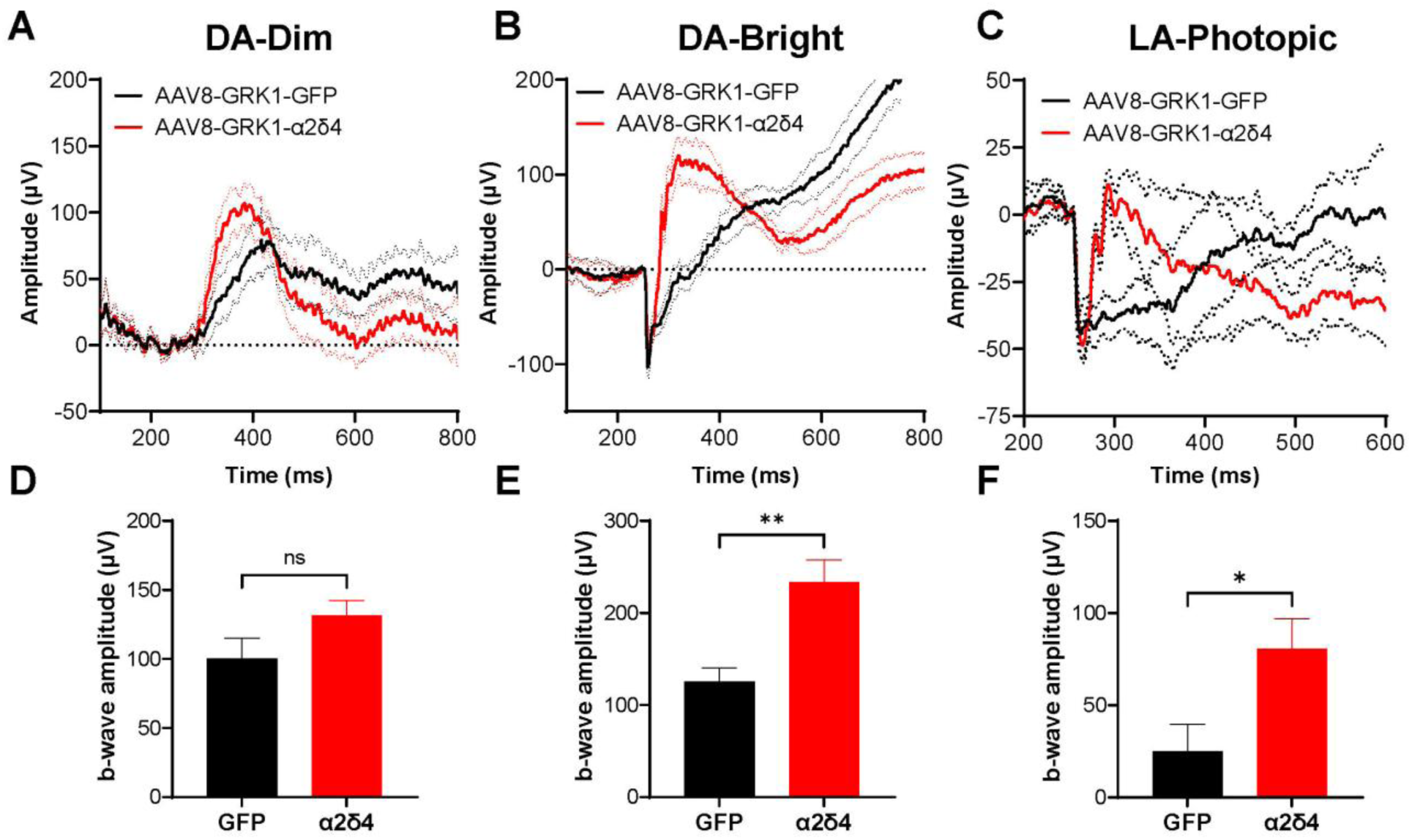
ERG studies demonstrate that rescue in middle aged α2δ4 KO (7-12M) restores synaptic function with reduced efficacy. **A-B.** Average traces of dark-adapted ERG response to light intensity of 2.479 (**A**), 6.316 (**B**) log photons/µm^2^ from rescued α2δ4 KO mice. **C.** Average traces of light-adapted ERG response to light intensity of 6.316 log photons/µm^2^ from rescued α2δ4 KO mice. **D-E.** Quantification of b-wave amplitudes of dark-adapted ERG responses at light intensities of 2.479 (**D**) and 6.316 (**E**) log photons/µm², in rescued α2δ4 KO mice (n = 5 mice). **F.** Quantification of b-wave amplitudes of light-adapted ERG responses at light intensities of 6.316 log photons/µm^2^ from α2δ4 KO mice (n = 5 mice). Error bars are SEM, unpaired t-test. *\*p < 0.05; **p < 0.01; points without an asterisk are not significant (ns)*.

At the structural level, reintroducing α2δ4 after 7 months of age still restored key features of photoreceptor synaptic organization. PSD95 and cone arrestin labeling were recovered within rod and cone terminals (Fig. 6A, B), although rod bipolar cell dendritic sprouting persisted and was not corrected by rescue (Fig. 6C). Similar to the young adult group, rescued middle-aged retinas showed restored synaptic ribbon labeling, Cav1.4 localization, and ELFN/mGluR6 organization in both the OPL and ectopic ONL regions (Fig. 6D, E).

**Figure 6.**
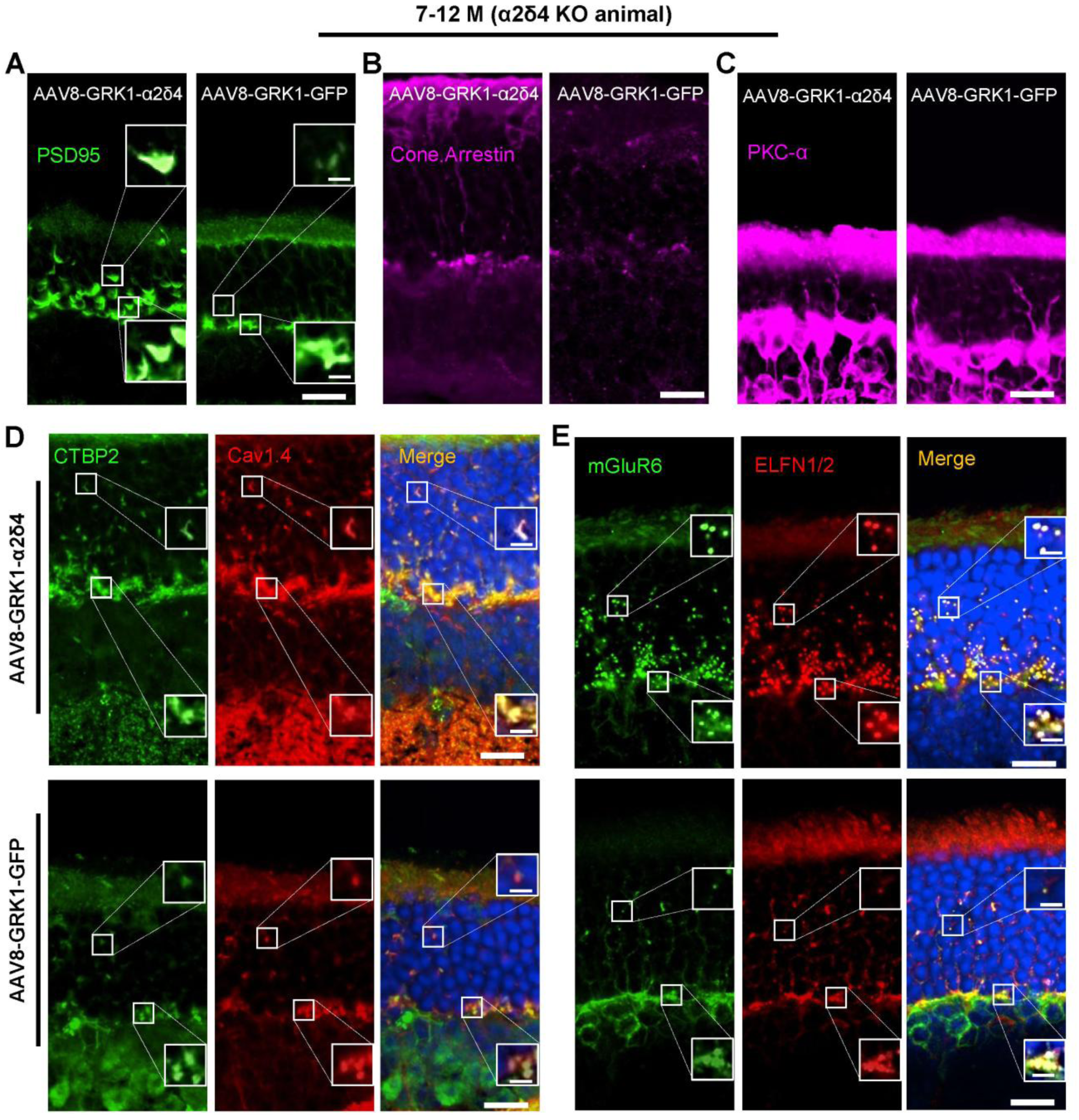
IHC studies demonstrate that rescue in middle aged α2δ4 KO (7-12M) restores synaptic assembly but not neurite remodeling. **A-C.** Representative confocal images of retinal sections from α2δ4 KO mice stained with anti-PSD95 (**A**), anti-cone arrestin (**B**), and anti-PKCα (**C**) to label photoreceptor terminals, cone photoreceptors, and rod bipolar cells (RBCs), respectively. Scale bar, 10 μm. Inset, 2.5 μm **D**. Representative confocal images of retinal sections from α2δ4 KO mice double immunolabeled for CTBP2 (green) and Cav1.4 (red). Scale bar, 10 μm. Inset, 2.5 μm **E**. Representative confocal images of retinal sections from α2δ4 KO mice double immunolabeled for mGluR6 (green) and ELFN1/2 (red). Scale bar, 10 μm. Inset, 2.5 μm

These data indicate that α2δ4 gene supplementation can restore synaptic molecular organization and improve visual pathway function even when delivered at a substantially later disease stage. However, the reduced rescue of dim-light scotopic responses suggests that rod circuit recovery is more sensitive to age than cone circuit recovery.

### The age-dependent functional rescue correlates with the spatial distribution of rescued synapses

The most prominent functional difference among age groups was observed under dim scotopic conditions. To identify potential factors underlying the age-dependent differences in functional rescue, we compared structural and functional measures across neonatal, young, and middle-aged adult rescue groups.

We first assessed viral transduction across age groups and confirmed that the AAV-infected retinal area was comparable among the three rescue cohorts (Suppl. Fig. 7A, B). Therefore, differences in rescue efficiency were unlikely to reflect major differences in viral delivery or transduction area.

To compare rescue efficiency across ages, we normalized b-wave amplitudes in each rescued eye to the GFP-treated control eye from the same animal and compared the fold increase among groups. Neonatal rescue produced the largest increase, followed by young adult rescue, whereas the 7- to 12-month group showed minimal increase which differed significantly from the neonatal group (Fig. 7A). In contrast, when rescue efficiency was evaluated using the normalized b/a ratio under bright dark-adapted or light-adapted conditions, the values were similar across the three age groups (Fig. 7B, C). These analyses further support that rod-dominant functional rescue is more age-sensitive than cone pathway rescue.

**Figure 7.**
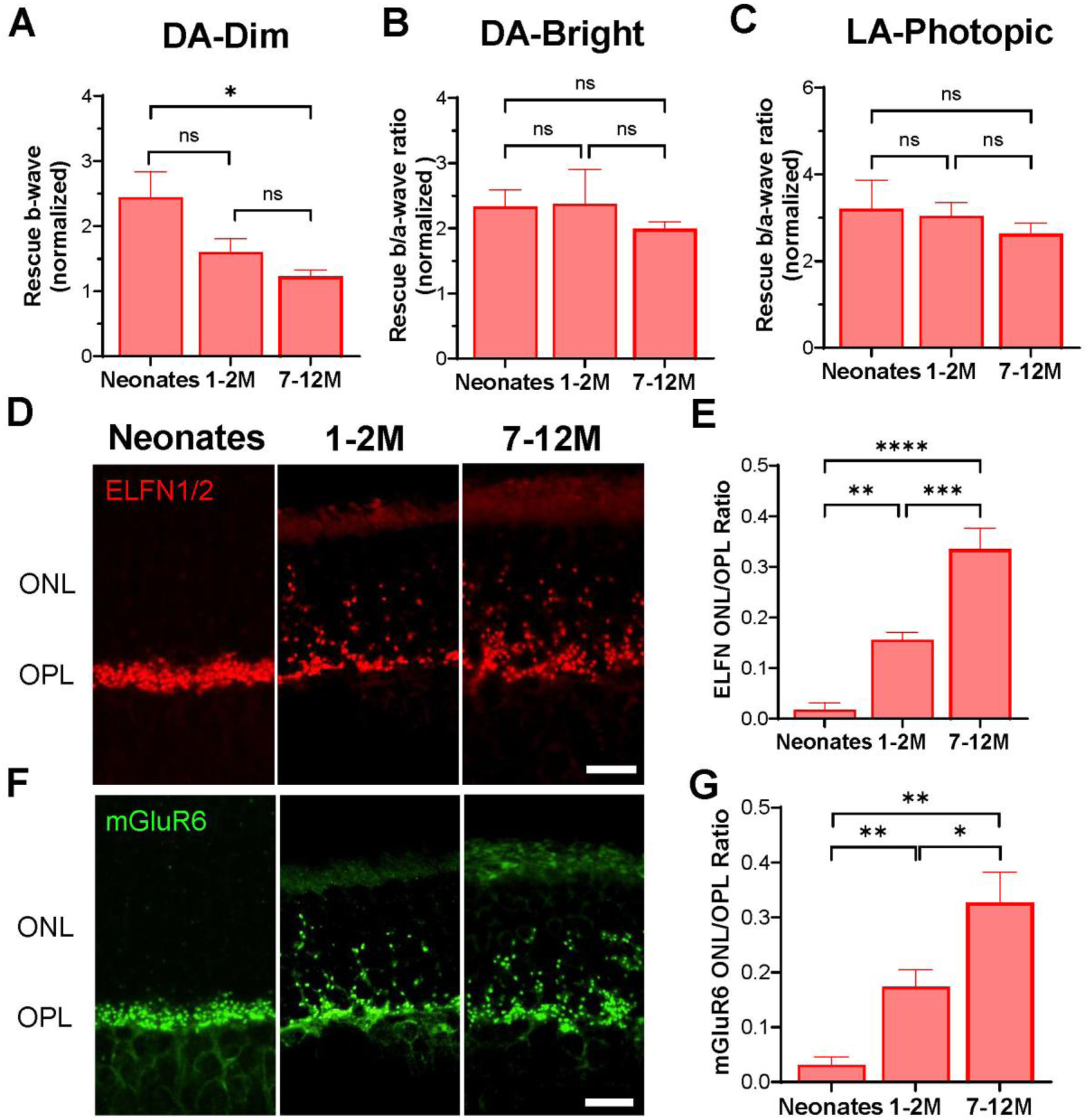
Age dependent functional rescue efficacy correlates with spatial distribution of rescued synapses. **A-C.** Normalized rescue b-wave in the dark-adapted ERG response at a light intensity of 2.479 log photons/µm² (**A**), and the normalized rescue b/a ratio under dark-adaption at 6.316 log photons/µm² (**B**) as well as under light-adaption at 6.316 log photons/µm² (**C**) in α2δ4 KO mice rescued at different time. **D-F** Representative confocal images of retinal sections from α2δ4 KO mice rescued at different times immunolabeled for ELFN1/2 (**D**) (red) and mGluR6 (**F**) (green). Scale bar, 10 μm. **E-G.** Quantification of the ELFN (**E**), and mGluR6 (**G**) density ratio (ONL/OPL) of α2δ4 KO retinas for different rescue groups (n = 3 mice per group). Error bars are SEM, unpaired t-test and one-way ANOVA followed by Tukey’s test. *\*p < 0.05; **p < 0.01, ***p < 0.01; points without an asterisk are not significant (ns)*.

Because synaptic markers ELFN and mGluR6 were robustly restored in all three rescue groups, we next asked whether the location of rescued synapses, rather than their molecular composition, contributed to differences in functional recovery. We quantified ELFN puncta density in rescued retinas across the ONL and OPL. Although total ELFN puncta density across retinal layers was comparable among groups (Suppl. Fig. 7C), synapse distribution differed markedly with age. Neonatal rescue produced the highest proportion of ELFN puncta in the OPL and the lowest proportion in the ONL, whereas the 7- to 12-month rescue group showed the opposite pattern, with more rescued puncta retained ectopically in the ONL (Fig. 7D, E; Suppl. Fig. 7D, E). Quantification of mGluR6 puncta showed a similar age-dependent localization shift from OPL to ONL (Fig. 7F, G; Suppl. Fig. 7F-H). Because most bipolar cell sprouting and most retracted photoreceptor terminals in α2δ4 KO retinas at both young and old age are rod bipolar cells and rod terminals, respectively ^21,31^, these ectopic ONL synapses are likely enriched for rod synapses ^21,31^. The parallel relationship between reduced OPL localization and diminished dim scotopic b-wave rescue suggests that ectopically localized ONL synapses, despite displaying organized molecular architecture, may support rod signaling less efficiently than synapses positioned within the OPL.

## Discussion

In this study, we tested whether AAV-mediated α2δ4 supplementation can restore photoreceptor synaptic structure and function in α2δ4-deficient retinas and whether rescue efficacy depends on the age at treatment. Our results demonstrate that α2δ4 re-expression is sufficient to reassemble key molecular components of both rod and cone synapses, including Cav1.4, ELFN proteins, and mGluR6, in neonatal, young adult, and middle-aged adult knockout retinas. However, the extent of functional recovery and correction of retinal remodeling differed substantially across ages and retinal circuits. Neonatal rescue restored synaptic molecular organization, normalized neurite positioning, and improved both rod- and cone-mediated function. In contrast, rescue in adult animals restored synaptic molecular organization and improved visual signaling but failed to reverse photoreceptor terminal retraction and bipolar cell dendritic sprouting (Fig. 8). These findings indicate that α2δ4-dependent synaptic assembly remains highly plastic even in mature retinas, whereas neurite positioning and rod circuit function are more constrained by developmental timing.

**Figure 8.**
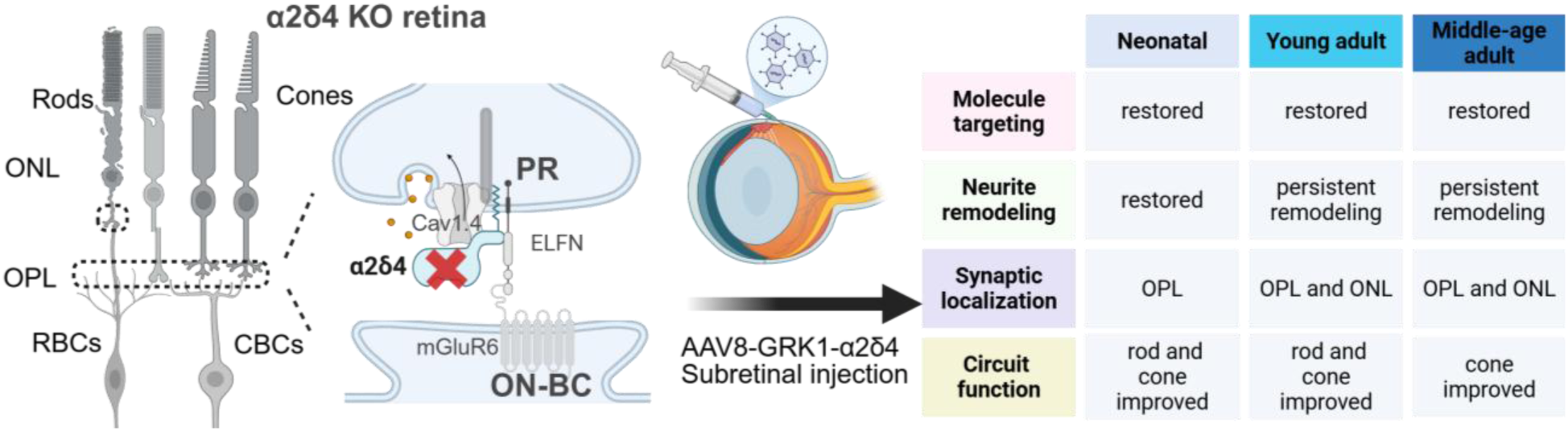
Schematic summarizing the age- and circuit-dependent AAV-α2δ4 rescue. Loss of α2δ4 leads to disrupted synaptic localization of Cav1.4, ELFN proteins and mGluR6, retraction of photoreceptor axons and sprouting of bipolar cell dendrites from OPL to ONL, and consequently, impaired circuit function. Gene supplement by AAV at different developmental stages achieved distinct rescue profiles shown in the table.

Our findings are consistent with a related preprint that became available during preparation of this manuscript ^32^. However, that study examined rescue at a single treatment time point and focused on rod-specific rescue, whereas our work demonstrates that therapeutic efficacy varies with both treatment age and circuit identity.

A major conclusion of this study is that rod and cone circuits exhibit distinct therapeutic windows for α2δ4 gene supplementation. Neonatal and young adult rescue significantly improved dim scotopic b-wave responses, whereas rescue in 7- to 12-month-old animals did not produce a significant improvement at rod-dominant scotopic flash intensities. In contrast, rescue under brighter dark-adapted conditions and under light-adapted conditions remained largely preserved across all treatment ages. Collectively, these findings suggest that cone circuits retain a broader therapeutic window than rod circuits, consistent with recent findings using a RP model ^33,34^. Although both rod and cone synapses require α2δ4 for normal organization and function, rod synapses are more vulnerable to age and other perturbations triggered remodeling while cone synapses are more resilient ^35–38^. This difference in plasticity could be one reason why AAV-α2δ4 is less capable of restoring rod circuit function after longstanding synaptic disruption.

Our quantitative analyses further support this idea by revealing that the progressive decline in dim scotopic rescue correlated with a shift in the spatial distribution of rescued synapses. This redistribution reflects the age-dependent circuit remodeling in α2δ4 KO retina ^31^ and provides a plausible explanation for the reduced recovery of dim scotopic responses in older animals. Thus, molecularly organized synapses can form outside the normal OPL environment, but their ability to support rod pathway transmission may be limited compared with synapses positioned within the appropriate retinal layer.

Several mechanisms may account for why ectopic synapse location reduces signaling efficiency despite restoration of molecular markers. First, displacement of photoreceptor terminals into the ONL may alter the geometry of the synaptic cleft, the apposition between glutamate release sites and postsynaptic mGluR6 complexes, or the extracellular space that shapes glutamate diffusion and clearance. Even subtle changes in cleft organization could affect the timing and concentration profile of glutamate reaching bipolar cell dendrites ^10,39,40^. Second, retraction of rod terminals may change the electrotonic relationship between the photoreceptor soma, inner segment, and synaptic terminal, potentially altering how voltage changes reach Cav1.4-containing active zones. Such changes could modify voltage propagation, calcium channel activation, or vesicle release efficiency ^41^, thereby limiting synaptic output even when Cav1.4 and associated synaptic proteins are restored. Third, ectopic synapses may lack layer-specific extracellular, glial, or regulatory cues that normally support efficient coupling between presynaptic release and postsynaptic GPCR signaling ^1,42–44^. These possibilities are not mutually exclusive and suggest that restoring synaptic molecular organization alone may not fully restore the biophysical and spatial context required for optimal rod transmission.

Another important finding is the dissociation between synaptic molecular organization and neurite remodeling, a distinction not shown in previous rescue studies of the Cav1.4 complex ^45,46^ and supported by the related preprint study ^32^. Across all rescue ages, α2δ4 expression restored Cav1.4, ELFN, and mGluR6 localization and generated synaptic puncta with similar molecular characteristics. In contrast, recovery of neurite positioning showed a strong dependence on developmental timing. Only neonatal rescue effectively corrected photoreceptor terminal retraction and rod bipolar cell sprouting, whereas adult rescue left substantial remodeling intact. These observations suggest that synaptic assembly and neurite elaboration are governed by distinct plasticity mechanisms ^47^. Synaptic organization appears relatively insensitive to the developmental stage or remodeling state of the neuron, provided that the critical synaptic proteins are present. In contrast, neurite positioning may depend on developmental programs that become progressively restricted with maturation and may involve calcium-dependent cytoskeletal mechanisms that are no longer fully accessible in adulthood given that animal model expressing non-conducting Cav1.4 exhibit similar neurite remodeling ^48^.

Our findings therefore highlight the potential importance of combining molecular rescue with strategies that stabilize or restore neurite architecture. Gene supplementation effectively reassembles synaptic protein complexes, but additional interventions targeting cytoskeletal regulation and related signaling pathways to stabilize neurites may be required to fully restore retinal circuitry ^49^. Such combinatorial approaches may be particularly important for diseases diagnosed after significant structural reorganization has already occurred.

In summary, AAV-mediated α2δ4 supplementation restores synaptic molecular organization in both developing and mature α2δ4-deficient retinas, demonstrating persistent plasticity of the photoreceptor synaptic assembly program. However, functional rescue is strongly influenced by both intervention time and circuit identity. Rod pathway rescue is most effective when treatment occurs early, whereas cone pathway rescue remains comparatively robust across ages. These findings define distinct therapeutic windows for synaptic assembly, neurite remodeling, and rod versus cone circuit recovery, and establish α2δ4 gene supplementation as a promising therapeutic strategy for inherited retinal synaptopathies caused by CACNA2D4 dysfunction.

## MATERIALS AND METHODS

### Antibodies

The following commercial antibodies were used: rabbit anti-cone arrestin (AB15282, Millipore), rabbit anti-Cav1.4 (365 003, Synaptic Systems), mouse anti-CTBP2 (612044, BD Biosciences), rabbit anti-ELFN1 (448 003, Synaptic Systems), mouse anti-PKCα (A31571, Invitrogen), rabbit anti-PSD95 (3450, Cell Signaling Technology), guinea pig anti-ribeye (192 104, Synaptic Systems), anti-α2δ4 OAAF04451,Avivasysbio), anti-mGluR6 (Cao 2011 ^50^) . Chicken anti-GFP (AB13970, Abcam).

### Adeno-associated virus (AAV)

Cloning of full-length mouse α2δ4 into a mammalian expression vector pcDNA3.1 was described previously ^21^. GRK1 promotor rAAV viruses (cAAV α2δ4 and GFP) were obtained from the Ocular Gene Therapy Core of University of Florida. The AAV8 vector is carrying a full-length mouse α2δ4 cDNA or GFP under the control of GRK1 to provide rod and cone specificity for α2δ4 expression ^28,29^.

### Animals

α2δ4 knockout (Wang et al., 2017) mice have been described previously. Both male and female mice were used in all experiments. Animals were housed under a controlled 12-hour light/12-hour dark cycle. Standard rodent chow and water were provided *ad libitum*, and mice were maintained in plastic cages with conventional bedding and enrichment. All animal procedures were conducted in accordance with National Institutes of Health guidelines and were approved by the University of Alabama at Birmingham Institutional Animal Care and Use Committee (IACUC).

### Immunohistochemistry

Eyes were enucleated, fixed in 4% paraformaldehyde for 15 minutes at room temperature, and cryoprotected overnight in 30% sucrose in PBS at 4 °C. Tissues were embedded in OCT compound (Fisher HealthCare, fisher scientific) and cryosectioned at 14 µm thickness. Sections were rehydrated in PBS, encircled with a hydrophobic barrier (PAP pen), and blocked for 1 hour at room temperature in PBS containing 0.1% Triton X-100 and 10% donkey serum. Primary antibodies were diluted in PBS containing 0.1% Triton X-100 and 2% donkey serum and applied for 1 hour at room temperature. Slides were washed 3–4 times for 5 minutes each in PBS with 0.1% Triton X-100.

Fluorophore-conjugated secondary antibodies (1:500 in PBS with 0.1% Triton X-100 and 2% donkey serum) were applied for 30 minutes at room temperature in the dark. Nuclei were counterstained with DAPI (DAPI Flouromount-G, SouthernBiotech) and coverslipped. Slides were cured overnight at room temperature and sealed with clear nail polish before storage at 4 °C. The antibodies used and their dilutions include: rabbit anti-Cav1.4, 1:100, rabbit anti-cone arrestin, 1:2000, mouse anti-CTBP2, 1:1000, rabbit anti-ELFN1, 5 µg/mL, sheep anti-mGluR6, 1:500, mouse anti-PKCα, 1: 250, rabbit anti-PKCα, 1:50000, mouse anti-PSD95, 1:5000, rabbit anti-PSD95, 1:100, guinea pig anti-ribeye, 1:500, rabbit anti-α2δ4, 1:250, chicken anti-GFP, 1:500.

### Confocal microscopy and image analysis

Images were acquired on a Nikon AX-R laser confocal microscope using 10×, 40×, and 60× objectives. The same acquisition settings were used for all experimental and control groups. Z-stacks were taken at 0.22 μm intervals, and maximum-intensity projections were generated using NIS Elements and Fiji (ImageJ).

#### mGluR6 and ELFN1/2 puncta

Regions of interest (ROIs) corresponding to the OPL and ONL were defined based on DAPI staining; the OPL ROIs were extended 2–3 rows of photoreceptor nuclei into the ONL, and the ONL ROIs were adjusted accordingly to remain non-overlapping. Cone pedicle ROIs were defined based on PNA lectin labeling. Within each ROI, mGluR6 and ELFN1/2 puncta were identified by Fiji’s Auto Threshold function (Moments algorithm), using background-subtracted (rolling ball radius = 5 pixels) maximum-intensity projections.

#### GFP Expression

The percent GFP expression was calculated by dividing the GFP-positive area of each wholemount, identified using Fiji’s Auto Threshold function (Yen algorithm), by the total wholemount area and multiplying by 100.

### Electroretinography (ERG)

Mice were dark-adapted overnight before recordings and anesthetized with ketamine/xylazine (80–100 mg/kg and 8–10 mg/kg, i.p.). Corneal anesthesia was achieved with topical 0.5% proparacaine, and pupils were dilated with 1% tropicamide and 2.5% phenylephrine. To maintain corneal hydration and facilitate electrode placement, a drop of 2.5% Hypromellose (Gonak, Akorn) was applied. ERGs were recorded from the left eye using a corneal contact lens electrode, with a gold wire loop electrode on the contralateral eye serving as the reference. Mice were placed on a temperature-controlled platform inside a Faraday cage, and light stimuli were delivered via a fiber optic positioned ∼1 cm from the corneal surface.

Dark-adapted (scotopic) ERGs were recorded in response to flash stimuli ranging from 1.527 log photons/µm² to 6.316 log photons/µm², followed by light-adapted (photopic) recordings after 5 minutes of rod-saturating background illumination. ERG signals were amplified, digitized, and acquired with a custom-built two-channel ERG system equipped with fiber optics to deliver flashes from either a 100-Watt halogen bulb or a high intensity LED source. A-wave amplitudes were measured from baseline to the trough, while b-wave amplitudes were measured from the trough of the a-wave to the subsequent peak. Implicit times were determined for both responses. Because KO traces lacked the characteristic b-wave peak, the implicit time of b-waves from eyes receiving AAV8-GRK1-α2δ4 was used as a reference to determine b-wave amplitudes in retinas from eyes receiving AAV8-GRK1-GFP. All responses were recorded and averaged using LabView (version 6.4.1, National Instruments), then exported into Microsoft Excel and analyzed in GraphPad Prism.

### Quantification and Statistical Analysis

Unpaired t-test and one-way ANOVA were used for all pairwise and multi-group comparisons as indicated in the figure legends, unless otherwise specified. Data are presented as mean ± SEM. Significance is indicated by \**p* ≤ 0.05, \*\**p* ≤ 0.01, \*\*\**p* ≤ 0.001, and \*\*\*\**p* ≤ 0.0001. Statistical analyses and graphing were performed using GraphPad Prism (Version 11.0.1)

## AUTHOR CONTRIBUTIONS

T.U. performed subretinal injection, ERG, IHC, confocal microscopy, collected and analyzed data, prepared figures, and contributed to manuscript writing; G.H. performed data analyses, image analysis, and contributed to manuscript writing; S.B. generated AAVs for subretinal injection; Y.W. designed the study, analyzed data, and wrote the manuscript.

## ACKNOWLEDGEMENTS

We wish to thank Dr. Choice Amieghemen for maintaining mice used in this study. We are grateful for the UF Vector Core for generating all the AAVs used for rescues study. We thank Dr. Jose Luis Roig-Lopez (UAB Vision Science P30 Core) for technical support. We thank Dr. Kirill Martemyanov (University of Miami) for sharing the α2δ4 KO mice and mGluR6 antibody. We also thank Dr. Tylor Lewis for critical reading of the manuscript. This work was supported by the National Institutes of Health/National Eye Institute (NIH/NEI) grant EY030554 (YW), Core Grant for Vision Research P30 EY003039, The E. Matilda Ziegler Foundation (YW), UAB IMPACT fund (YW), and the UAB Vision Science Research Center pilot grant (YW). Microsoft Co-Pilot was used to edit the manuscript.

## CONFLICT-OF-INTEREST STATEMENT

The authors declare no competing interests.

## Abbreviations

AAV: adeno-associated virus
BC: bipolar cells
CBC: cone bipolar cell
CSNB: congenital stationary night blindness
CTBP2: C-terminal-binding protein 2
ELFN1: extracellular leucine rich repeat and fibronectin type III domain containing 1
ELFN2: extracellular leucine rich repeat and fibronectin type III domain containing 2
ERG: electroretinography
GFP: green fluorescent protein
GRK1: G protein-coupled receptor kinase 1
IHC: immunohistochemistry
IRDs: inherited retinal diseases
KO: knockout
MGLUR6: metabotropic glutamate receptor 6
ONL: outer nuclear layer
OPL: outer plexiform layer
PKC-α: protein kinase C alpha
PSD95: postsynaptic density protein 95
RBC: rod bipolar cell**ROIs:** regions of interest
VGCC: voltage-gated calcium channel
WT: wild type

## Notes

### Competing Interest Statement

The authors have declared no competing interest.

