## supplement material for "AAV-mediated age- and circuit-dependent restoration of photoreceptor synaptic structure and function in α2δ4-associated retinal synaptopathy"

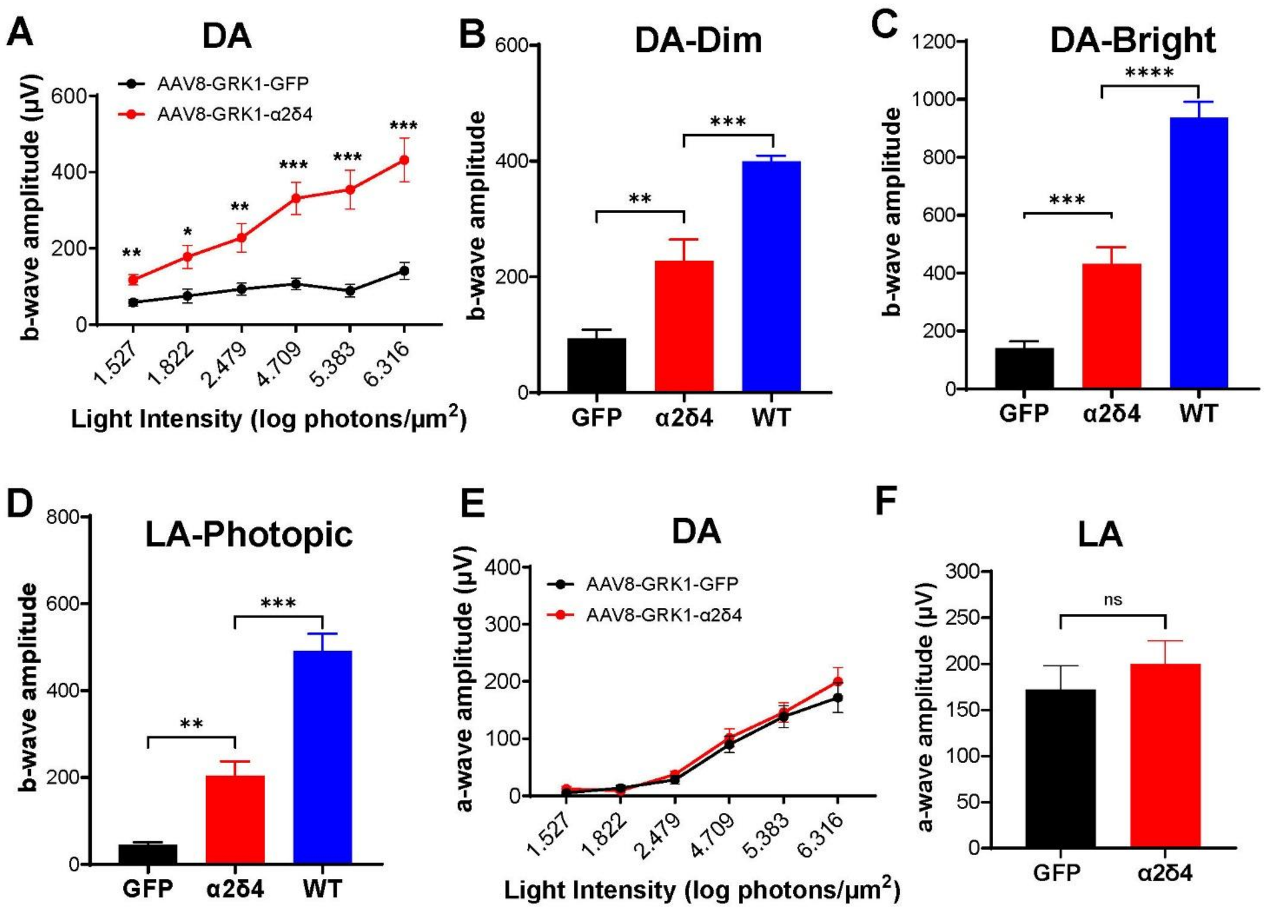


**Supplementary Figure 1 (related to Figure 1). AAV8-GRK1-α2δ4 rescue in α2δ4 KO neonates improves ERG responses.**

**A, E.** Quantification of b-wave amplitudes (**A**), and a-wave amplitudes (**E**) across increasing light intensities in α2δ4 KO mice (n = 11 mice).

**B-C.** Quantification of b-wave amplitudes of dark-adapted ERG responses at light intensities of 2.479 (**B**) 6.316 (**C**) log photons/µm2, which stimulate both rods and cones from α2δ4 KO and non-injected WT mice (n = 11 for KO mice, n=6 for WT).

**D.** Quantification of b-wave amplitudes of light-adapted ERG responses at a light intensity of 6.316 log photons/µm2, which stimulate cones from α2δ4 KO and non-injected WT mice (n = 9 mice for KO and 6 for WT).

**F**. Quantification of a-wave amplitudes at a light intensity of 6.316 log photons/µm2 in α2δ4 KO mice (n = 11 mice).

Error bars are SEM, multiple unpaired t-test (**A, E**) and unpaired t-test (**B, C, D, D**). **p < 0.05; **p < 0.01, ***p < 0.001; ****p < 0.0001 ns; not significant.*

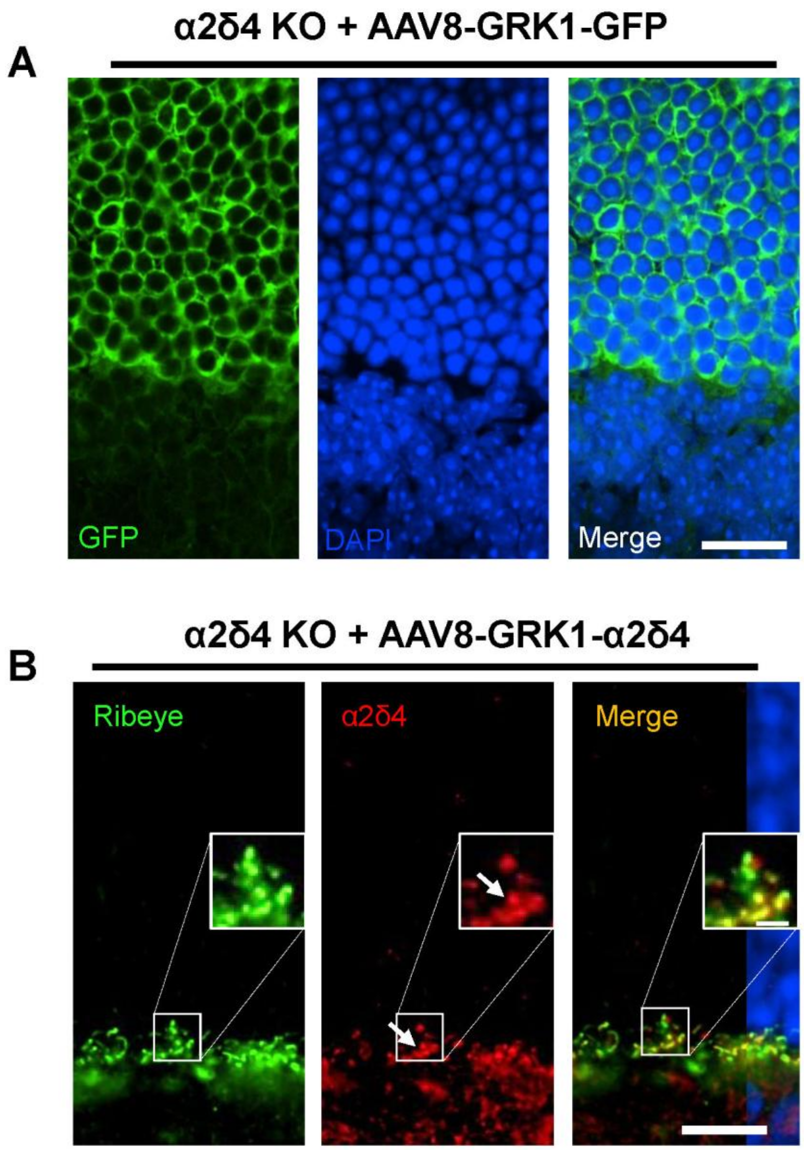


**Supplementary Figure 2 (related to Figure 2): IHC studies demonstrate that AAV-mediated α2δ4 expression in both rod and cone synapses of KO retinas.**

**A.** Representative confocal images of retinal sections from treated α2δ4 KO mice immunolabeled for GFP (green) and DAPI staining (blue) showing GFP is specifically expressed in the photoreceptor layer. Scale bar, 10 µm.

**B.** Representative confocal images of retinal sections from treated α2δ4 KO mice double immunolabeled for Ribeye (green) and α2δ4 (red) showing correctly targeted α2δ4 in both rod and cone (white arrow) synapses. Scale bar, 10 µm. Inset, 2.5 µm.

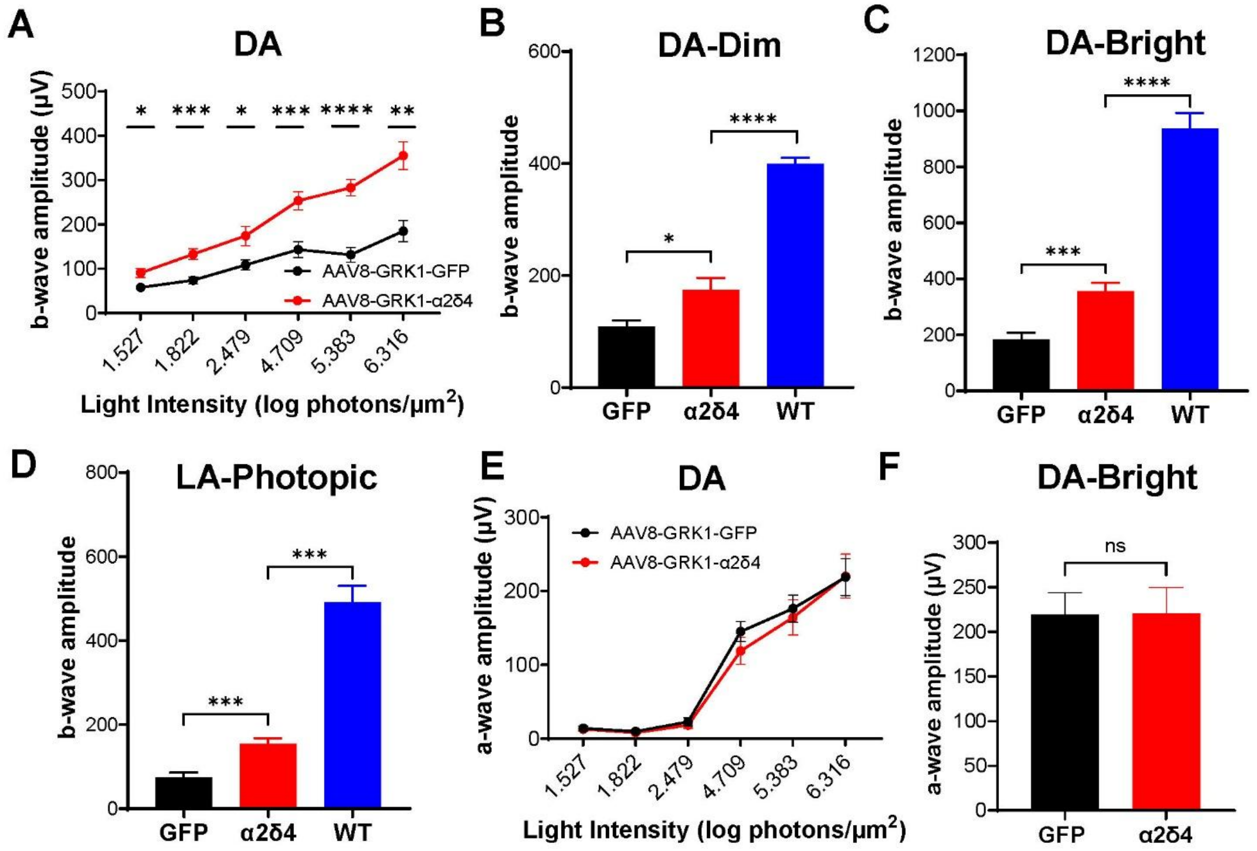


**Supplementary Figure 3 (related to Figure 3): AAV8-GRK1-α2δ4 rescue in α2δ4 KO 1-2M improves ERG responses.**

**A, E.** Quantification of b-wave amplitudes (**A**), and a-wave amplitudes (**E**) across increasing light intensities in α2δ4 KO mice (n = 13 mice).

**B-C.** Quantification of b-wave amplitudes of dark-adapted ERG responses at light intensities of 2.479 (**B**) 6.316 (**C**) log photons/µm2, which stimulate both rods and cones from α2δ4 KO mice (n = 13 mice for KO and 6 mice for WT).

**D.** Quantification of b-wave amplitudes of light-adapted ERG responses at a light intensity of 6.316 log photons/µm2, which stimulate cones from α2δ4 KO mice (n = 12 mice for KO and 6 mice for WT).

**F**. Quantification of a-wave amplitudes at a light intensity of 6.316 log photons/µm2 in α2δ4 KO mice (n = 13 mice).

Error bars are SEM, multiple unpaired t-test (**A, E**), one-way ANOVA (**B, C, D)** and unpaired t-test **(F**). **p < 0.05; **p < 0.01, ***p < 0.001; ****p < 0.0001 ns; not significant.*

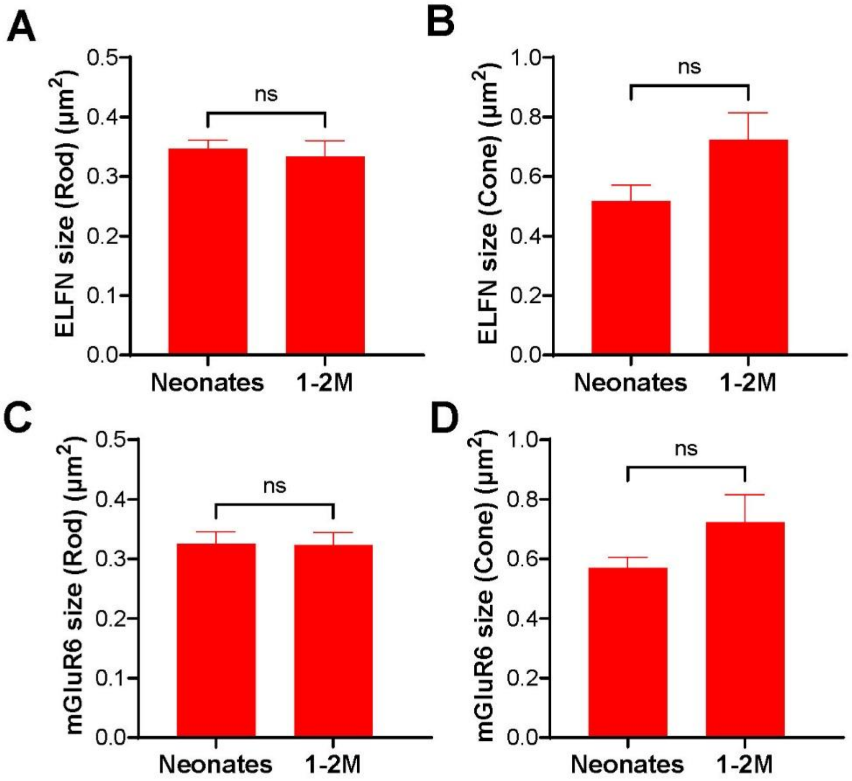


**Supplementary Figure 4 (related to Figure 4): Quantitative analysis of rescued synaptic size in neonatal and 1-2M old α2δ4 KO mice.**

**A-B.** Quantification of the ELFN puncta size in rod (**A**) and cone synapses (**B**) of rescued α2δ4 KO retinas (n = 3 mice per each group).

**C-D.** Quantification of the mGluR6 puncta size in rod (**C**) and cone synapses (**D**) of rescued α2δ4 KO retinas (n = 3 mice per each group).

Error bars are SEM, unpaired t-test. *ns; not significant.*

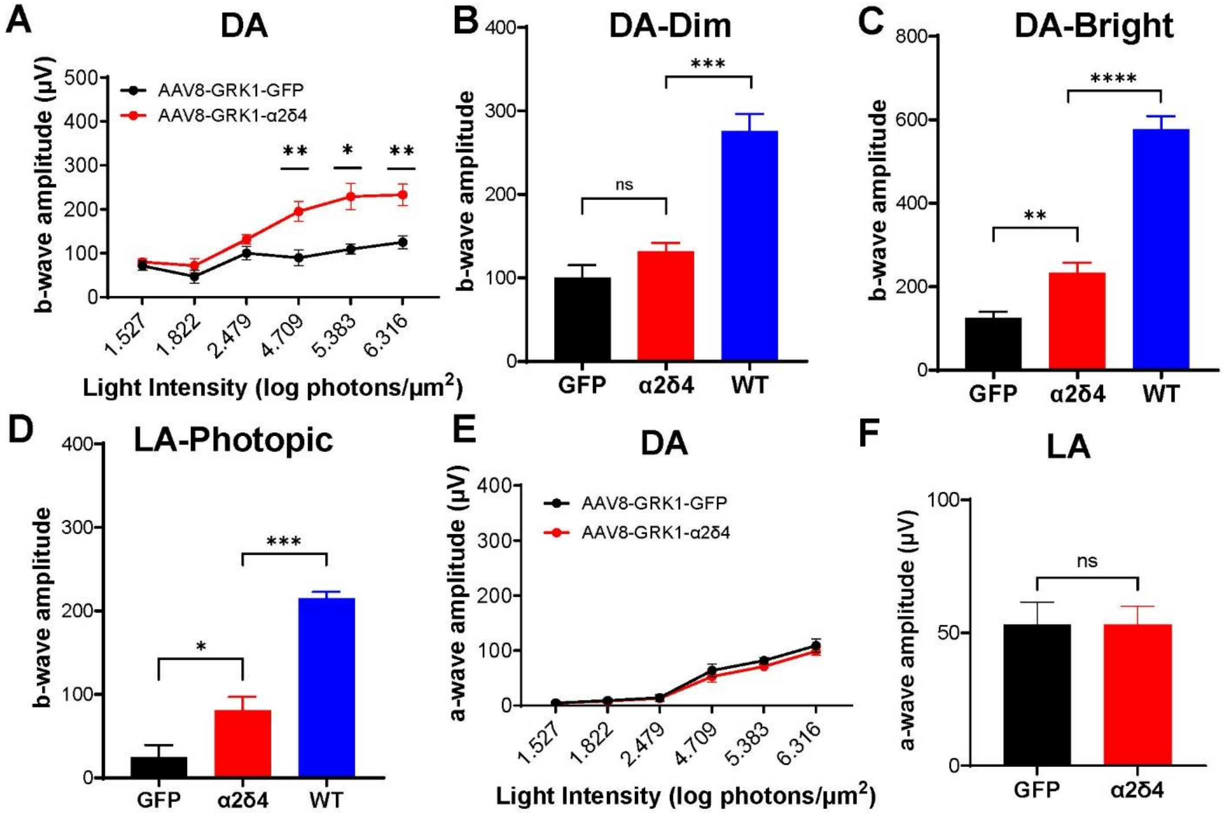


**Supplementary Figure 5 (related to Figure 5): AAV8-GRK1-α2δ4 rescue in α2δ4 KO 7-12M improves cone-mediated ERG responses.**

**A, E.** Quantification of dark-adapted (DA) b-wave amplitudes (**A**), and a-wave amplitudes (**E**) across increasing light intensities in α2δ4 KO mice (n = 5 mice).

**B-C.** Quantification of b-wave amplitudes of dark-adapted ERG responses at light intensities of 2.479 (**B**) 6.316 (**C**) log photons/μm2, from α2δ4 KO (n = 5 mice) and non-injected WT (n= 5 mice) animals.

**D.** Quantification of b-wave amplitudes of light-adapted (LA) ERG responses at a light intensity of 6.316 log photons/μm2, from α2δ4 KO (n = 5 mice) and non-injected WT (n= 5 mice) animals..

**F**. Quantification of light-adapted ERG a-wave amplitudes at a light intensity of 6.316 log photons/μm2 in α2δ4 KO neonates (n = 5 mice).

Error bars are SEM, multiple unpaired t-test (**A, E**), one-way ANOVA (**B, C, D**) and unpaired t-test (**F**). **p < 0.05; **p < 0.01, ***p < 0.001; ****p < 0.0001 ns; not significant.*

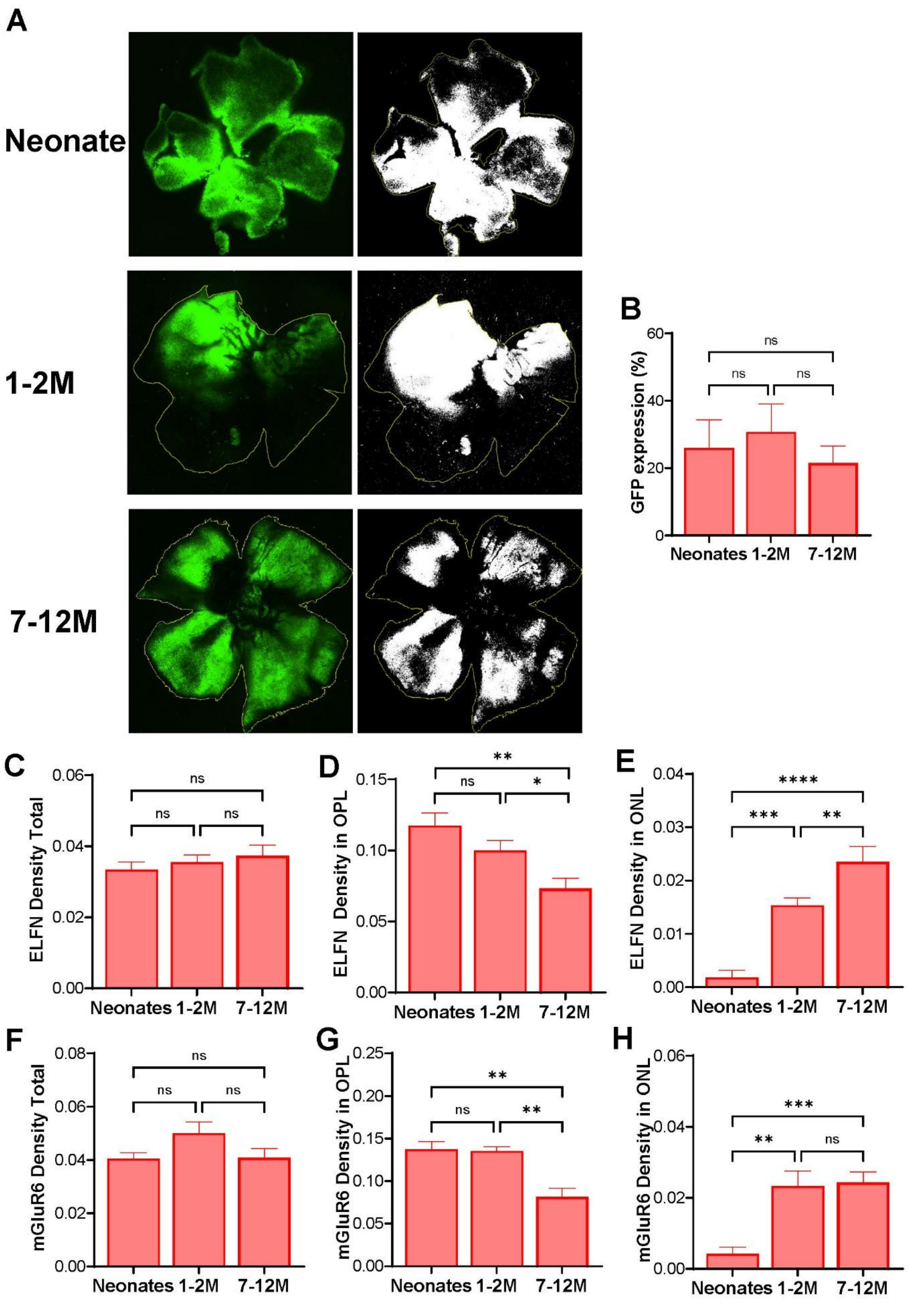
A
Neonate
1-2M
7-12M
B
GFP expression (%)
ns
ns
ns
Neonates 1-2M 7-12M
C
ELFN Density Total
ns
ns
ns
D
ELFN Density in OPL
**
ns
*
E
ELFN Density in ONL
****
***
**
F
mGluR6 Density Total
ns
ns
ns
G
mGluR6 Density in OPL
**
ns
**
H
mGluR6 Density in ONL
***
**
ns
Neonates 1-2M 7-12M

**Supplementary Figure 6 (related to Figure 7). Comparative analysis of the spatial distribution of rescued synapses across neonate, young and mid-aged adult rescue groups.**

**A.** Representative retinal flat mount shows robust GFP expression in neonates (up-left panel), 1-2M (middle-left panel), and 7-12M (low-left panel) rescue animals.

**B.** Quantification and comparison of GFP expression (%) in retinal whole mounts across groups.

**C-E.** Quantification of total ELFN puncta density (**C**), its density in OPL (**D**), as well as in ONL (**E**).

**F-H.** Quantification of total GluR6 puncta density (**F**), its density in OPL (**G**), as well as in ONL (**H**).

Error bars are SEM, unpaired t-test and one-way ANOVA. $*p < 0.05$; $**p < 0.01$, $***p < 0.001$, $****p < 0.0001$, *ns; not significant.*
